# Long read sequencing reveals sequential complex rearrangements driven by Hepatitis B virus integration

**DOI:** 10.1101/2021.12.09.471697

**Authors:** Songbo Wang, Jiadong Lin, Xiaofei Yang, Zihang Li, Xu Tun, Tingjie Wang, Bo Wang, Liangshuo Hu, Kai Ye

**Author notes:** Correspondence to Kai Ye.

## Abstract

Integration of Hepatitis B virus (HBV) into human genome disrupts genetic structures and cellular functions. Here, we conducted multiplatform long read sequencing on two cell lines and five clinical samples of HBV-infected hepatocellular carcinomas (HCC). We resolved three types of viral integration-induced complex genome rearrangements (CGR) and proposed a model of ‘multi-hits and sequential-breaks’ to depict their formation process by differentiating inserted HBV copies with HiFi long reads. We deduced that all three complex types were initialized from focal replacement and fragile virus-human junctions triggered subsequent rearrangements. We further revealed that such rearrangements caused a prevalent loss-of-heterozygosity at chr4q, accounting for 19.5% of HCC samples in ICGC cohort and contributing to immune and metabolic dysfunction. Overall, our long read based analysis reveals novel sequential rearrangement processes initiated by HBV integration, hinting its structural and functional impact on HCC.

---

Persistent hepatitis B virus (HBV) infection is reported to be responsible for >50% of hepatocellular carcinoma (HCC) worldwide^1,2^. Integration of HBV fragments into human genome happens in more than half of HBV-infected HCC (36 out of 61)^3^, causing genome instability and onco-gene expression alternation, eventually leading to carcinogenesis^4^.

Several studies utilized short read sequencing (SRS) to survey genomic and oncogenic preference of HBV integrations sites^5–7^. However, due to short read-length (typically 100-200bp), SRS detects genomic preference of human-virus junctions but fails to reveal complexity of whole events. With long read sequencing (LRS) such as CLR and ONT, HBV integration events have been classified into either intra- or inter-chromosome alterations, either directly affecting cancer driver genes such as TERT and MYC^1,8^ or inducing focal gain, lost and dicentric chromosome by translocations^1,9,10^. Successful as they are to link viral integration and human complex genome rearrangements (CGR), long reads have not provided us a detailed survey of virus integration events due to high error rates of CLR or ONT and median coverage data (less than 20X)^9,10^. We expect that deep LRS especially accurate HiFi sequencing would be ideal to characterize complex structure of inserted HBV DNAs10 and their underlying formation mechanisms

In this study, we applied multi-platform deep LRS technologies, such as CLR, ONT and CCS, on 7 HBV-infected HCC (HBV-HCC) samples, consisting of 2 cell lines (SNU-182 and SNU-387) and 5 clinical samples **(Supplementary Table 1)**. Collectively, we provided a panaroma for four types of HBV integrations, in total 16 events **(Online methods, Fig. 1a, Fig. 1b and Supplementary Fig. 1)**. Type I and II were characterized as focal alterations caused by viral integration. Type III involved complex and integration-induced unbalanced translocations while Type IV referred to the most complex event, chromothripsis. Specifically, after characterization of numbers of ‘hits’ (integrations) and rounds of ‘breaks’ (double strand breaks, DSB) for the observed four types, we proposed a Multi-hIts and Sequential-breaKs (MISK for short) model to describe formation processes of those integration-induced CGRs from initial integration state (**Fig. 1c)**. Finally, we utilized HCC WGS and RNA-seq data from ICGC cohort to explore the prevalence of our discoveries and their implications on immune and metabolic functions.

**Fig. 1.**
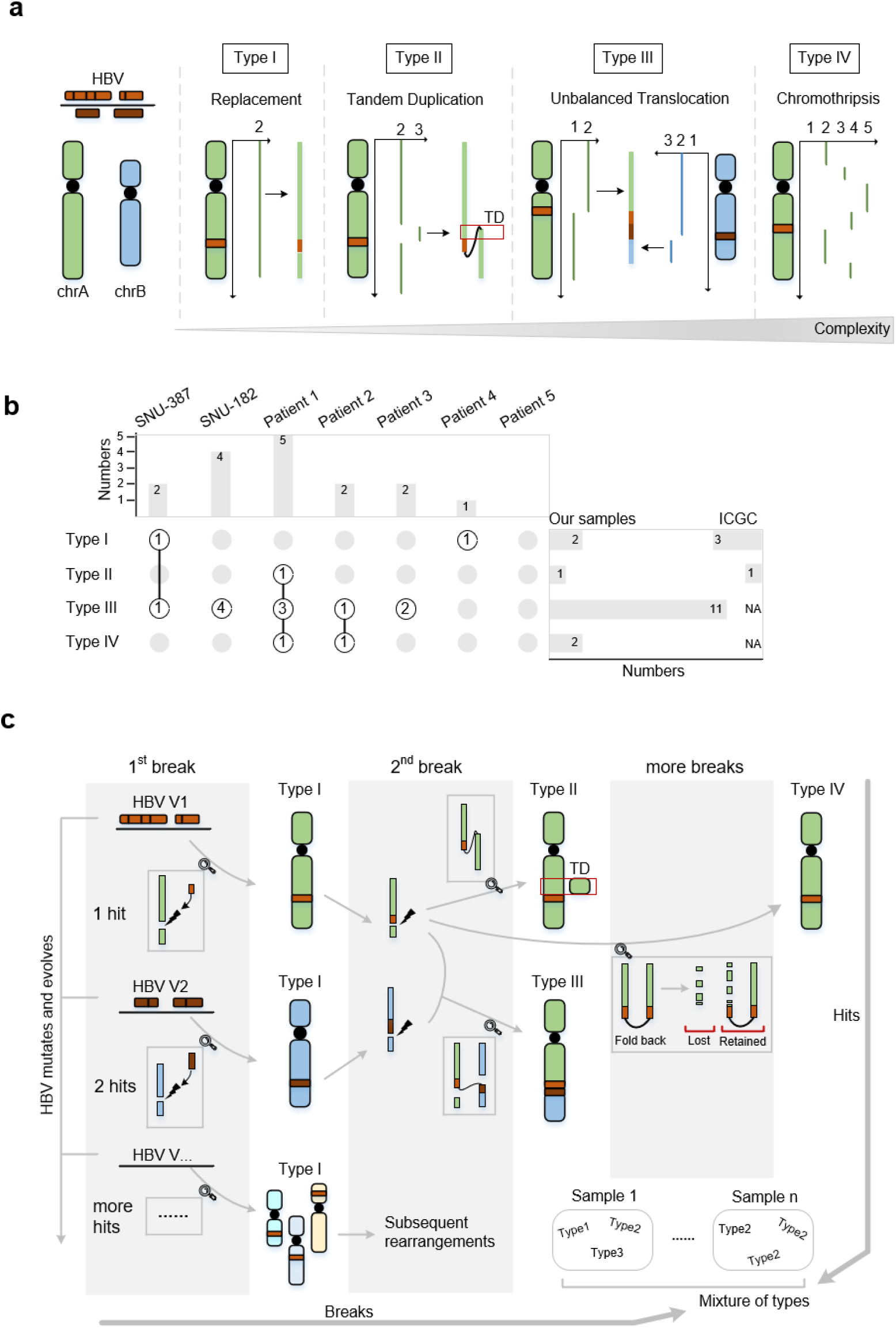
Introduction of novel discoveries.

In the Type I and II integrations, HBV hits human genome once and causes focal alterations by either replacing short human sequence (n=2) or tandemly duplicating genes (n=1) with two HBV fragments, as evidenced by single LRS read threading through human-virus-human junctions (**Fig. 2a and 2c**). Based on the features observed in LRS data, we deduced three Type I and one Type II focal alterations from published single human-virus junctions in ICGC HCC dataset^3^ **(Online methods, Supplementary Figs. 2 and 3)**. Previous study has reported promoter- or exon preferred integrations at *TERT* or *KMT2B*^1^. Similarly, tandem duplications directly affected genes, doubling copies of cancer-related genes such as *KCTD14* and *FSIP2* **(Fig. 2c and Supplementary Fig. 3)**. Replacements were gene-related as well. For instance, two HBV fragments replaced about 12bp of intron sequence of *CDH1* (chr16:68,744,185-68,744,197) in Event16 **(Fig.2a)**.

**Fig 2.**
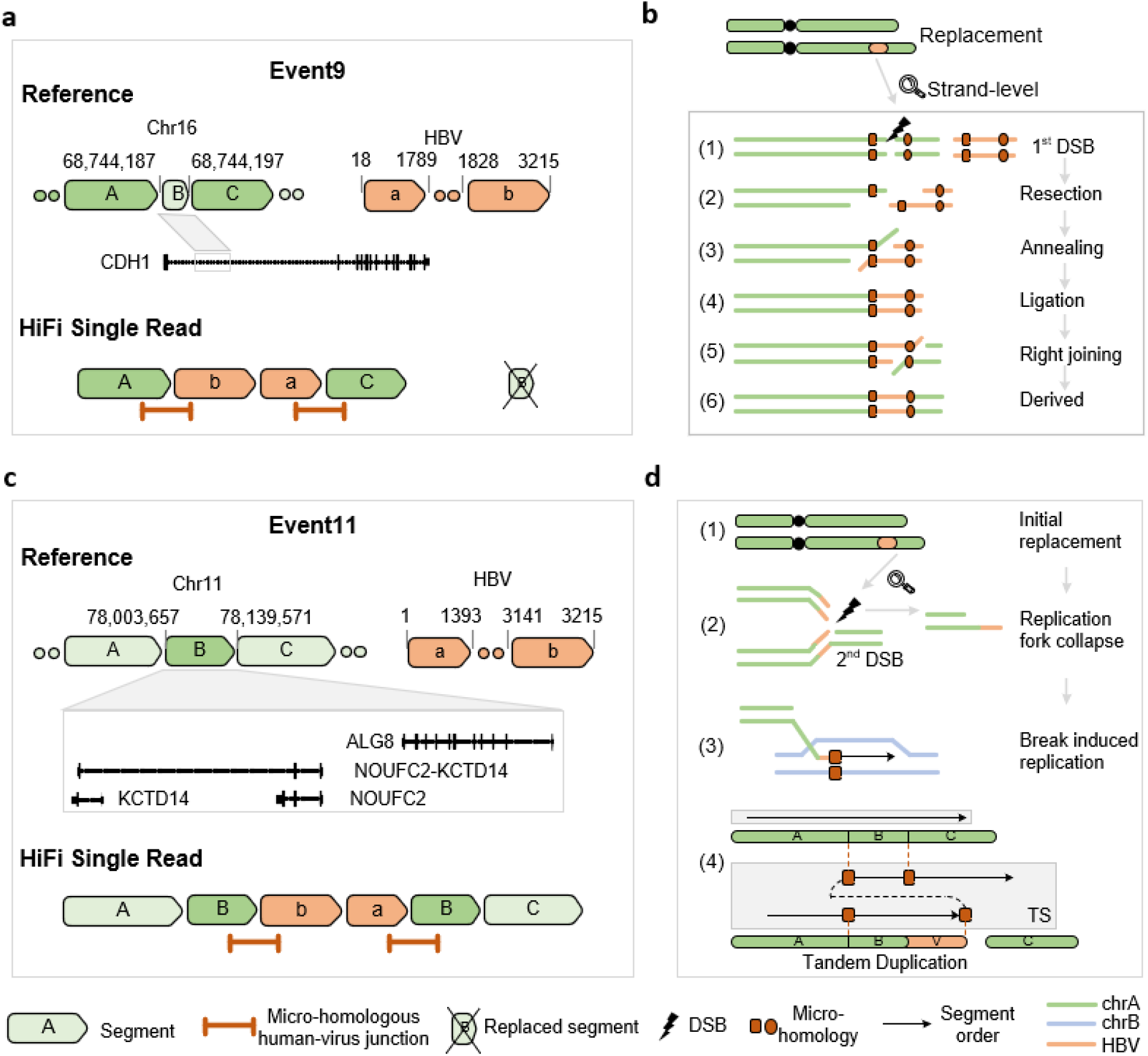
Type I and Type II.

The short replaced sequence and micro-homologies flanking two human-virus junctions supported a known mechanism mediating HBV integration^11^, the micro-homology mediated end-joining (MMEJ) repair pathway, often leading to short deletions right at DSB^12^. We considered the formation of the replaced sequence as the consequence of micro-homologies annealing and flap trimming during MMEJ **(Fig.2b)**. Those ‘one-hit and one-break’ replacements, were usually benign, but initiated instability of genome, especially at the human-virus junctions. As a result, the initial step of our MISK model was accomplished and ready for subsequent rearrangements through a second round of DSB followed by more complicated DNA repair pathways, including micro-homology mediated break-induced replication (MMBIR)^13^, mediating the progress from replacements to tandem duplications by homology-dependent template switching (TS, **Fig.2d**).

The Type III complex events (n=11, **Supplementary Fig. 4**) caused large scale copy number variations (CNVs) through unbalanced-translocations. As a result, one or more integrated HBV fragments bridged two chromosomes together. Both two virus-human junctions linking two chromosomes were right at the boundaries of significantly altered CNAs, characteristic with one gain (GOH) and one loss (LOH) of heterozygosity (**Fig. 3a)**. Among those unbalanced translocations, a ‘two-hits and two-breaks’ process characterized six of them as evidenced by multiple distinct inserted HBV copies. Facilitated by HiFi long reads in SNU-182, we could distinguish HBV strains by building a phylogeny tree with precisely detected single-nucleotide-variants (SNVs, **Online Methods, Supplementary Fig. 5**). For instance, in Event3, chromosome 4 and 8 were bridged together by two distinct strains of HBV (**Fig. 3a),** harboring characteristic SNVs and locating at non-adjacent branches in the phylogeny tree. **(Fig. 3b)**.

**Fig 3.**
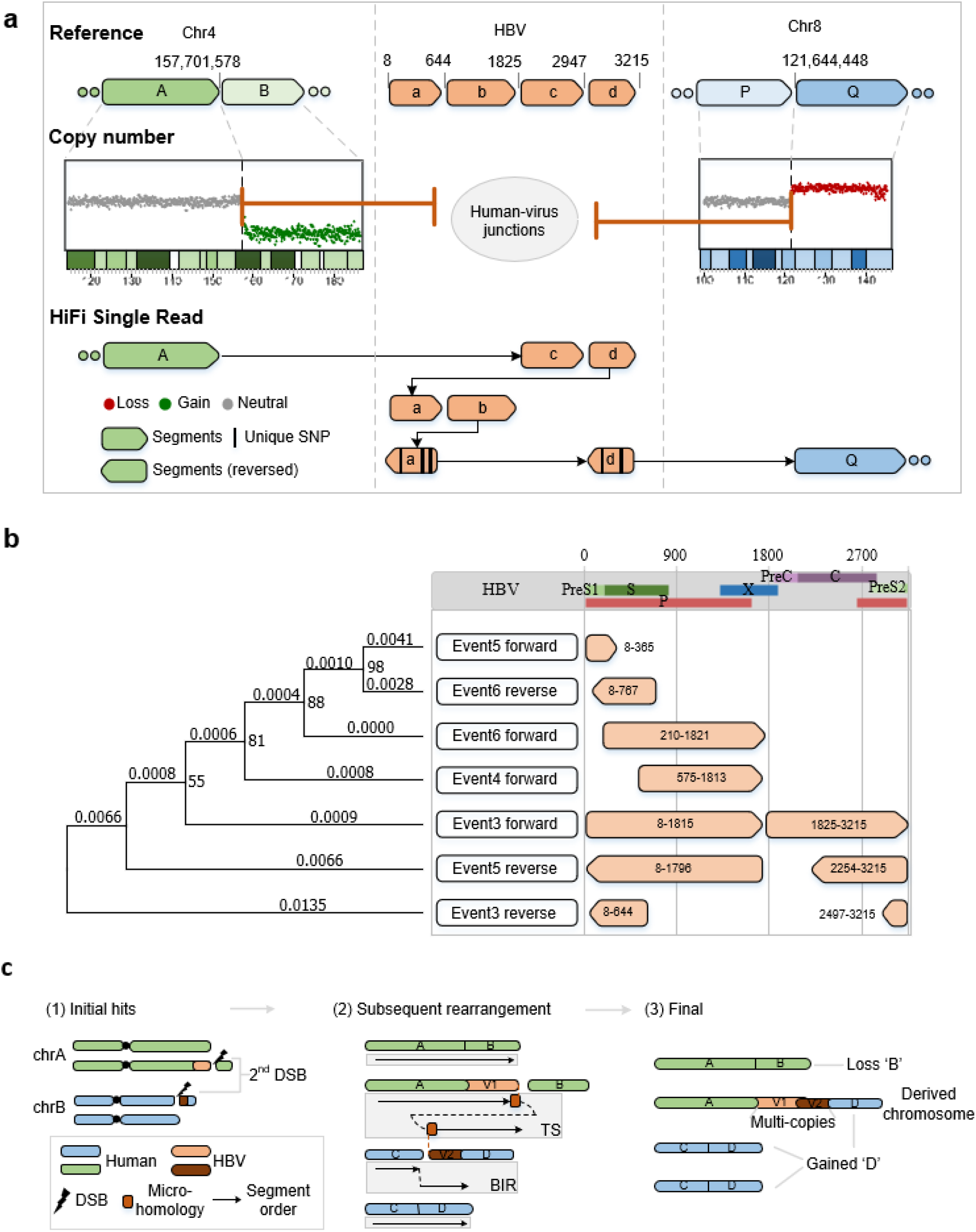
Type III.

The proposed MISK model well depicted the sequential formation process of the ‘two-hits and two-breaks’. **(Fig. 3c)**. We argue that during persistent infection, HBV hit two different chromosomes at different time points (two-hits) through first round DSBs (1^st^-DSB) mediated by MMEJ^11^, yielding two Type I replacement events (marked (1) in **Fig. 3c**). Later, fragile human-virus junctions broke and exposed DSBs again (2^nd^-DSB, two-breaks). Finally, via MMBIR^13^, the replication template switched from one DSB to the other via micro-homologies between two HBV copies and an unbalanced translocation formed eventually (marked (2) in **Fig. 3c**).

The rest of five unbalanced translocations contained only one copy of HBV DNA **(Supplementary Fig. 6)**. However, CNVs and micro-homologies identified between virus-human junctions still supported MMBIR after the initial MMEJ-mediated integration. We deduced the above as ‘one-hit and two-breaks’ **(Supplementary Fig. 7)**, where after the 2^nd^-DSB, the template switched from one DSB, caused by virus-human junctions, to another DSB occurring on human genome.

The most complex Type IV (n=2, **Supplementary Fig. 8 and 9**), integration-induced chromothripsis, was identified from clinical samples in this study. For instance, in Event 12, we observed chromosome-wide CNVs and 30 genomic rearrangements localized between chr14 and chr16 **(Methods and Fig. 4a)**. Most of those rearrangements located right at the boundaries of copy number neutral, loss or gain regions, implying that chr14 and chr16 underwent a chromothripsis event. Moreover, HBV integrated into the boundary of the rightmost amplified region and formed a symmetrical fold-back-inversion structure (**Supplementary Fig. 8**). An additional fold-back-inversion structure lay right at the boundary of the highest amplified region **(Fig. 4a)**. These two nested fold-back-inversion structures indicated breakage-fusion-bridge (BFB) repair^14^, a known mechanism to trigger chromothripsis^15^.

**Fig. 4.**
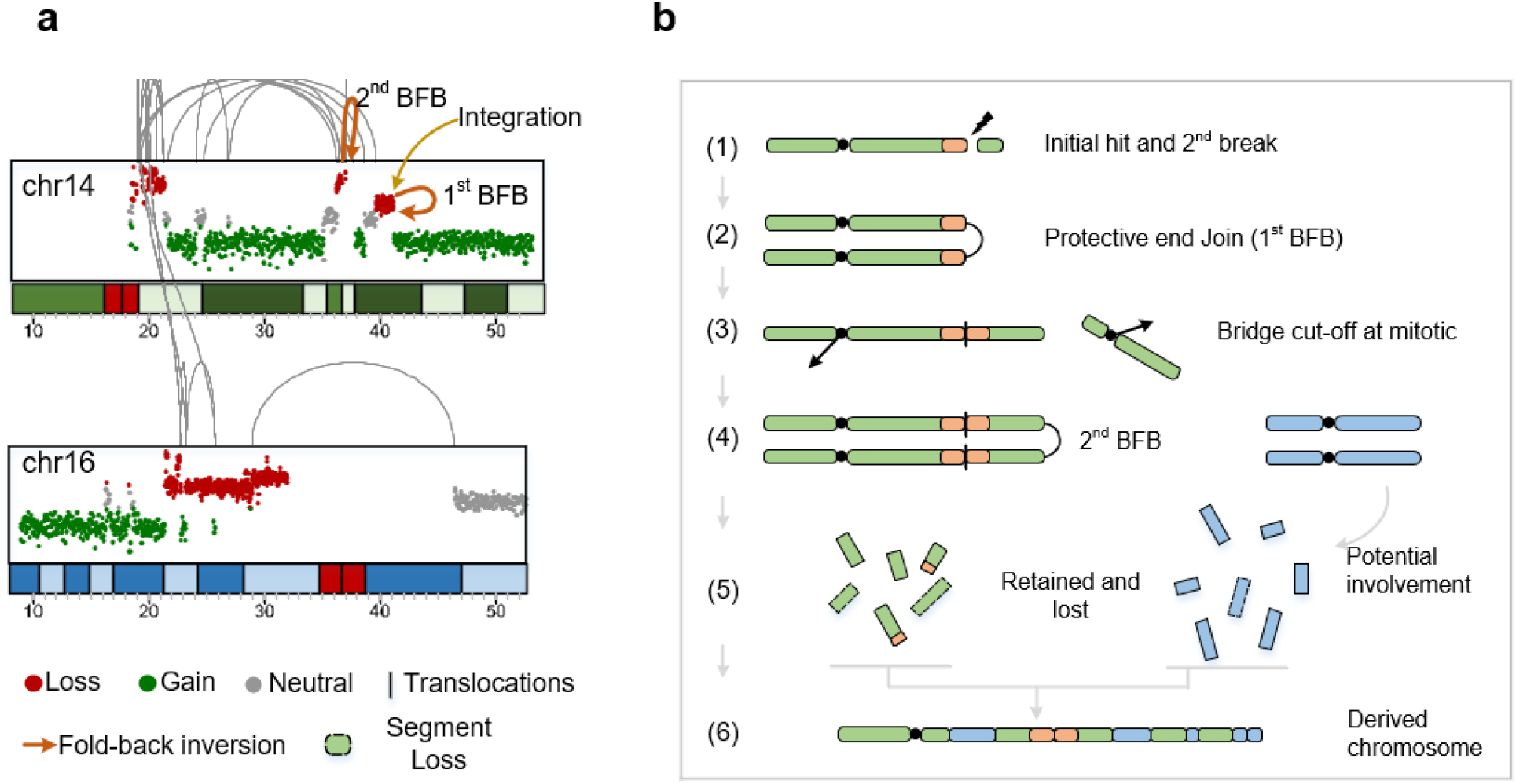
Type IV.

Given the previously proposed tenets^16^, we argued that the two observed chromothripsis events were actually subsequent rearrangements of initial HBV integration through a ‘one-hit and multi-breaks’ process (**Fig. 4b, Supplementary Note**). Specifically, HBV first integrated into chr14 as Type I replacement (one-hit, 1^st^-break, marked (1) in **Fig. 4b**). Secondly, at the integration site, telomeric DSB happened (2^nd^-break) and the unprotected ends was fused by non-homologous end-joining (NHEJ) during DNA replication (marked (2) in **Fig. 4b**). Later, the formed dicentric chromosome was cut off during mitosis and the second BFB happened in the next cell replication cycle, eventually leading to chromothripsis due to local DNA fragmentation produced by bridge breakage^15^ (multi-breaks).

The proposed four types could **concur** or **occur repeatedly** in a single HCC sample **(Fig. 1b, Supplementary Table 1)**. We investigated the four events in cell line SNU-182 and built a phylogeny tree built with all HBV copies detected in HiFi-sequenced SNU-182 (Fig. 3b). We observed that seven HBV copies from four events scattered at different branches and nodes in the tree, indicating that different initial Type I integrations probably hit the genome at different time points during chronic infection, supporting the proposed MISK model. Accompanied with rounds of breaks, these initial hits were rearranged to form more complex Type II, III or IV events, leading to various combinations of four integrations types in HCC samples (Fig. 1c).

As proposed in MISK model, subsequent rearrangements occurring after the initial HBV integrations, often led to severe CNVs on human genome. Notably, in the seven samples of this study, we observed a frequent LOH at chr4’s long arm (chr4q-LOH for short) in five samples, three of which were caused by Type III events. It has been reported that chr4q-LOH presented in 11% of ovarian cancer^17^. We examined 303 HCC samples from ICGC cohort (**Supplementary Table 2**) and found that ~19.5% of them (n=59) harbored chr4q-LOH **(Supplementary Table 3)**. In addition, if HBV infection was required (HBV-HCC), chr4q-LOH prevailed in ~32% samples, while ~12% in NBNC-HCCs, suggesting HBV infection as a significant high risk of chr4q-LOH (Chi-square Test, *P*=0.003). Please note that ICGC HCC samples were sequenced with SRS technology, probably leading to much fewer integration sites (5/59)^9^ while LRS in this study revealed much higher proportion (3/5) of HBV integrations in chr4q-LOH breakpoints.

Next, we examined the hotspots of these chr4q-LOH breakpoints in 64 samples (5 from our samples, 59 from ICGC) and found two hotspot breakpoint clusters among 53 of them **(Fig. 5a, Supplementary Table 3)**. Cluster1 (n=16) concentrated at chr4q33-34, leading to LOH at chr4q35, where HCC suppressor genes, *IRF2* and *ING2*^18^, were lost but without significant expression alteration. However, two immune associated genes, *ACSL1* and *F11*^19–21^, also located at chr4q35 and were induced significantly down-regulated (Kruskal-Wallis Test, *P*=0.0012 and 0.0035, respectively, **Methods and Fig. 5b)**. The CN loss of *ACSL1* and *F11*, together with another 14 down-regulated genes were enriched in KEGG item complement and coagulation cascades, indicating an aberrant immune system. (*P.adj*= 6.13E-08, **Fig. 5d, Supplementary Tables 4 and 5)**.

**Figure 5.**
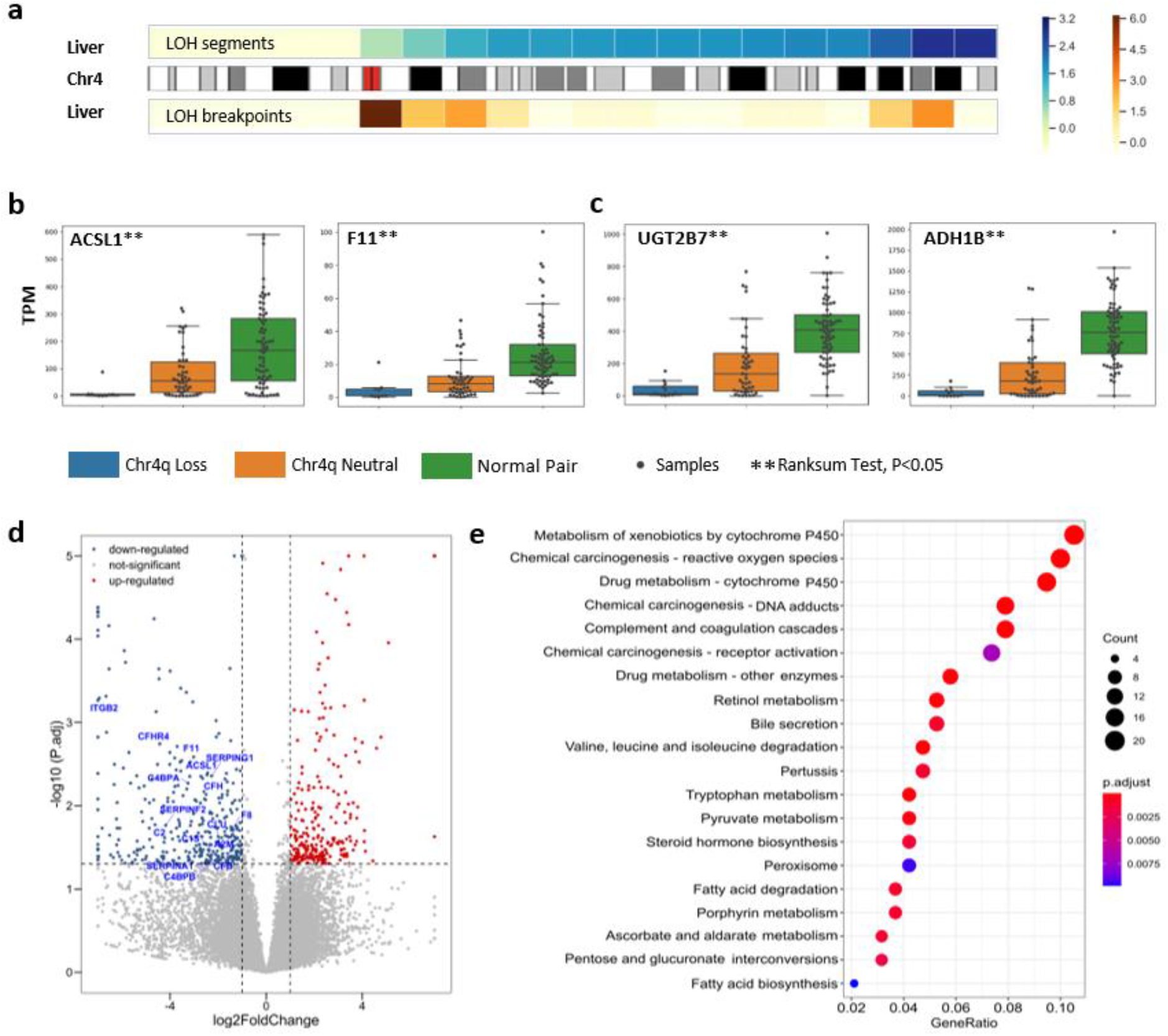

In addition to chr4q35-LOH (Cluster1), we discovered another enriched cluster at chr4q12-13 region, leading a much larger LOH from chr4q13 to 3’ telomere (37 out of 59, Cluster2). Besides the two immune-related genes mentioned above, metabolic genes at the downstream of chr4q13, such as *UGT2B7*^22^ and *ADH1B*^23^, were significantly down-regulated **(Fig. 5c)**. As a consequence, 22 metabolism pathways were significantly enriched **(Supplementary Fig 9, Supplementary Table 6)**, suggesting deficient metabolism functions induced by chr4q-LOH.

Those functional analysis suggested that although random HBV integrations led to various copy number changes^24^[], the copy number loss of specific regions, such as chr4q12-13 and chr4q33-34, could cause immune and metabolic dysfunction, leading to HCC. We argue that majority of initial random HBV integrations might be benign as the proportion of genomic regions harboring exons and promoters is rather small (12 out of 109 integration sites^3^), but subsequent rearrangements cause deleterious alterations of gene copies and expressions, leading to HCC.

In this study, based on characterization of seven HBV-HCC samples with deep LRS, we proposed the MISK Model to illustrate the sequential formation processes of integration-induced CGRs with involvements of multiple DNA repair mechanisms such as MMEJ, MMBIR and BFB. Moreover, we showed two dominating clusters of LOH at chr4q in HCC, mostly attributed to the Type III unbalanced translocations, leading to aberrant immune and metabolism.

LRS allows us to characterize the full-picture of HBV integration. However, LRS is still too short to resolve any chromothripsis events and manual reconstructions are required to integrate CNs and translocations. Current HBV integration studies are conducted on the end-state HCC where the landscape of integration status among early stages of HCC is lacking. In this study we deduced the MISK model based on both inter-sample and intra-sample comparison of HBV integration events identified from the end-state HCC samples. The basic principle of the MISK model is ‘integration then rearrangement’, where integration-induced CGRs were a subsequent rearrangement process from initial replacement. However, ‘integration when rearrangement’ is also possible if MMEJ-mediated HBV integration and MMBIR-mediated unbalanced translocations could take effect at the same cell division phase (**Supplementary Note**, **Supplementary Fig. 10**). Therefore, genome sequencing of samples that undergo different infection and carcinoma stages towards HCC will be certainly beneficial for our understanding of the role HBV playing in HCC.

HBV integrations have been discussed and explored with SRS and LRS in the past decade. We demonstrate that LRS especially the HiFi LRS can comprehensively characterize the complex integration patterns, mechanisms and impacts. Our work lays down an important foundation for further large-scale HBV integration studies with LRS, hinting new strategy for future diagnose and treatment.

## Online Methods

### Cell lines

SNU-182 and SNU-387 were acquired from the National Collection of Authenticated Cell Cultures. The SNU-387 were sequenced on the PacBio Sequel machine, producing the CLR reads while SNU-182 were sequenced on the PacBio Sequel II sequencer, generating HiFi reads. Briefly, libraries for single molecule real-time (SMRT) PacBio genome sequencing were constructed following the standard protocols. The high molecule genomic DNA was sheared to ~20Kbp targeted size, followed by damage repair and end repair, blunt-end adaptor ligation and size selection. The average CLR sequencing coverage was 45 reads per bp while that forThe average HiFi sequencing coverage was 24 reads per bp.

Additionally, the RNA sequencing of SNU-182 and SNU-387 were produced. A total amount of 1 g RNA per sample was used for library preparation, and the library was generated using NEBNext UltraTM RNA Library Prep Kit for Illumina (NEB, USA), following the manufacturer’s recommendations. Then, the prepared library was sequenced on the Illumina Novaseq platform, generating the 150bp paired-end reads.

### Clinical samples

Five primary HCC samples were collected from the First Affiliated Hospital of Xi’an Jiaotong University. Five collected tumor tissues were sequenced on the Nanopore PremethION sequencer according to the manufactures’ instructions. In general, the long DNA fragments were qualified and selected using the PippinHT system (Sage Science, USA), and the A-ligation reaction were conducted with NEBNext Ultra II End Repair/dA-tailing Kit. The adapter in the SQK-LSK109 (Oxford Nanopore Technologies, UK) was used for further ligation reaction and DNA library was measured by Qubit 4.0 Fluorometer (Invitrogen, USA). Finally, about 700ng DNA library was constructed for each tumor tissue and sequenced with PremethION. The average ONT sequencing coverage was 45 reads per bp.

### ICGC cohort

Project LIRI-JP and LIHC-US were selected from ICGC cohort. Filtering out those in black list, we total obtained 303 HCC samples with WGS, 118 of which have RNA-seq. The corresponding normal paired data were also collected. We utilized the corresponding profiles of CNA and transcripts read counts aggregated by PCAWG working group. The viral condition for each sample was also obtained from previous studies as well^3,25,26^.

#### Characterization of HBV integration with LRS

HBV integration pattern characterization was performed by a self-developed pipeline, including: (1) read mapping (2) candidate reads selection; (3) cluster reads to events; and (4) classification of integration events

1. Read mapping. Raw reads from the LRS technologies were aligned to GRCh38 with HBV genome via Minimap2^27^. The preset parameter of Minimap2, ‘map-pb’ and ‘map-ont’ were activated for PacBio and Oxford Nanopore (ONT) reads, respectively. Besides soft-clip and MD tag were set by – Y and −MD. Parameter −a and --eqx was set to output SAM format with =/X CIGAR operators. Then we used SAMtools^28^ to sort and index the output.
2. Candidate reads selection. We selected reads that span both human and viral genomes by traversing reads that partially aligned on HBV genome. Minimum map quality was set to 20 to filter imprecise reads. For each candidate read *r_i_*, we collect all its primary and supplementary alignments and sorted them by read coordinates. Therefore, *r_i_* can be represented as all its alignments: *r_i_* =< (chr1, start1, end1, T), (HBV, start2, end2, F),… (chr3, start2, end2, T) >, where T and F denoted the mapping direction.
3. Cluster reads to events. We cluster candidate reads to integration events. Different reads but with similar alignments were merged together based on their coordinates. We set the maximum difference between two alignments to 50bp. When comparing two reads, we considered their orientations and toke the minus strand into account. After merging, integration events with less than 5 supporting reads were filtered out. We totally obtained 17 integration events, 3 of which had centromere alignments and were excluded.
4. Classification of integration events. Integration events were classified to 3 types, including focal replacement, unbalanced translocation and chromothripsis. Reads that supporting focal replacement usually included 4 alignments: the middle two aligned to HBV while the left and right segments aligned focally and closely on the same chromosome. The unbalanced translocations featured as at least 3 alignments, two of which aligned to different chromosomes and the left aligned to HBV. The chromothripsis always included fold back inversions formed by HBV fragments. As a result, we could observed a nearly symmetrical arrangement of alignments. Accompanying with the CNA and inter-/intra-translocations, we determined the chromothripsis event.

#### Defer HBV integration pattern from NGS samples

Since SRS could not fully cover the whole event because the limitations of short read length, we deferred Type I focal replacements and tandem duplications from single human-virus junctions in PCAWG’s previous study []. First, in each sample, we clustered each two junctions with same chromosome and close coordinates (distance on human genome was required less than 100Kb), acting as a potential focal event. Then, we acquired already aligned (hg19) WGS BAM file and profiled its sequence coverage using IGVtools with zoom level (-z) as 7 and window size (-w) as 1000. Finally, given the junctions and coverage profile, if the coverage between two virus-human junctions was lower than that on the other side, we considered it as a replacement event. Similarly, if the coverage was doubled between the two junctions, we considered it as a tandem duplication event.

#### Phylogeny tree

We used HiFi sequenced SNU-182 to build a phylogeny tree. Firstly, we separated apart the aligned HBV fragments by the four events. We also separated the different oriented segments in one event. Because HBV always fragmented and some sequences might lost, we then fixed the gap region with reference genome sequences. As a result, we obtained 7 individual HBV mutated genomes. We used MEGA-X^29^ (GUI mode) to build a neighbor-joining tree using maximum composite likelihood method. Both transitions and transversions of substitutions were included. The test of phylogeny ran bootstrap method 500 replications.

#### Copy number variation analysis

For LRS, genome-wide profile of CNA was performed by CNVkit^30^ without paired control samples. We ran CNVkit under the default settings except the bin size was set to 100Kbp, and copy number plots were created with the scatter option. In addition, we applied the IGVtools^31^ to all chromosomes at 100Kbp resolution, generating the binned whole genome coverage. The binned coverage was saved as TDF format for further visualization. For SRS (ICGC samples), CNV profiles of all samples were obtained from PWAWG working group. The published results included total CN, major CN and minor CN of each chromosome segment. We further implemented a simple script to visualize the results for better understanding.

LOH or GOH in ICGC samples were determined by relative change between two neighboring CN segments. CN segments with length less that 10Kbp were excluded to avoid sequence coverage bias. To avoid aneuploidies which were common in cancer, we strictly required that the CN change must start from neutral CN (CN=2). Therefore, LOH meant CN change from 2 to 1 while GOH meant CN change from 2 to 3. Besides, we required LOH or GOH to extend to the telomere. Samples whose chr4 satisfied the above conditions were kept and the corresponding change breakpoints were kept as well. Chr4-neutral samples’ chr4 kept CN=2.

#### Chromothripsis detection

Chromothripsis featured as massive CNVs and inter-/intra-chromosome translocations. CNV breakpoints of our samples were obtained from CNVkit mentioned above. Translocation detection for chromothripsis followed a similar pipeline as that for HBV integration. We collected all abnormal reads with two alignments aligned in different chromosomes (inter-translocation) or in same chromosome but aligned two distant region (> 1Mb, intra-translocation). After clustering single reads and filtering by minimum supporting reads (default 5), we assigned each translocation breakpoint with CNV breakpoint by imitating the coordinate’s differences within 50bp. Those inter-/intra-translocations without CNV were excluded. Finally, we visualize CNV and translocations together utilizing karyoploteR^32^.

#### Gene expression and pathway enrichment

ICGC HCC samples were contributed into 4 groups. Group1 and group2 were samples from chr4q-LOH breakpoint Custer1 and Cluter2, respectively. Group3 included all chr4-netural HCC samples. Group4 was the set of all normal-paired samples of group 1, 2 and 3. We performed two rounds of analysis: <group1, group3> and <group2, group3> to reveal the impaction of different chr4q-LOH clusters on genes. Besides, we also performed <group3, group4> as a control to understand the gene expression alteration after tumorigenesis. Gene transcript read count matrix and TPM matrix was obtained from PWAWG working group^33^. Differential transcript expression between each two groups was analyzed by DESeq2^34^. Genes with P.adjust < 0.05 were considered as significantly altered. Besides, genes with log2FoldChange > 1 or <-1 were set to ‘UP’ and ‘DOWN’, respectively for further pathway enrichment. For genes with more than one transcript, we chose the most significant one and removed the left. The visualization (volcano plot) was done by R package ggpubr (https://rpkgs.datanovia.com/ggpubr). KEGG and Go pathway enrichments were perform with clusterProfiler^35^. Up- and down-regulated genes were enriched separately. The p and q value cutoff were both set to 0.05. Pathway items that satisfied the p and q value cutoff and had more than 10 genes enriched were kept and visualize by R package enrichplot (https://yulab-smu.top/biomedical-knowledge-mining-book/).

## Acknowledgements

K.Y. and X.Y. are supported by the National Science Foundation of China (32125009, 32070663, and 62172325), the Key Construction Program of the National ‘985’ Project, the World-Class Universities (Disciplines), and the Characteristic Development Guidance Funds for the Central Universities and the Fundamental Research Funds for the Central Universities.

## Author contributions

K.Y. designed and supervised research. S.W. and J.L performed HBV integration analysis. S.W., T.X. and T.W performed the gene expression analysis. B.W. and L.H. performed clinical surgeries and offered the clinical HCC samples. Z.L. performed Sanger analysis of HBV-human junctions. K.Y., S.W., J.L. and X.Y. wrote the paper. All authors read and proved the final version of this manuscript.

## Competing interesting

The authors declare no competing interests.

## Notes

### Competing Interest Statement

The authors have declared no competing interest.

