## Supplementart_figs for "Long read sequencing reveals sequential complex rearrangements driven by Hepatitis B virus integration"

#### Supplementary Figure 1

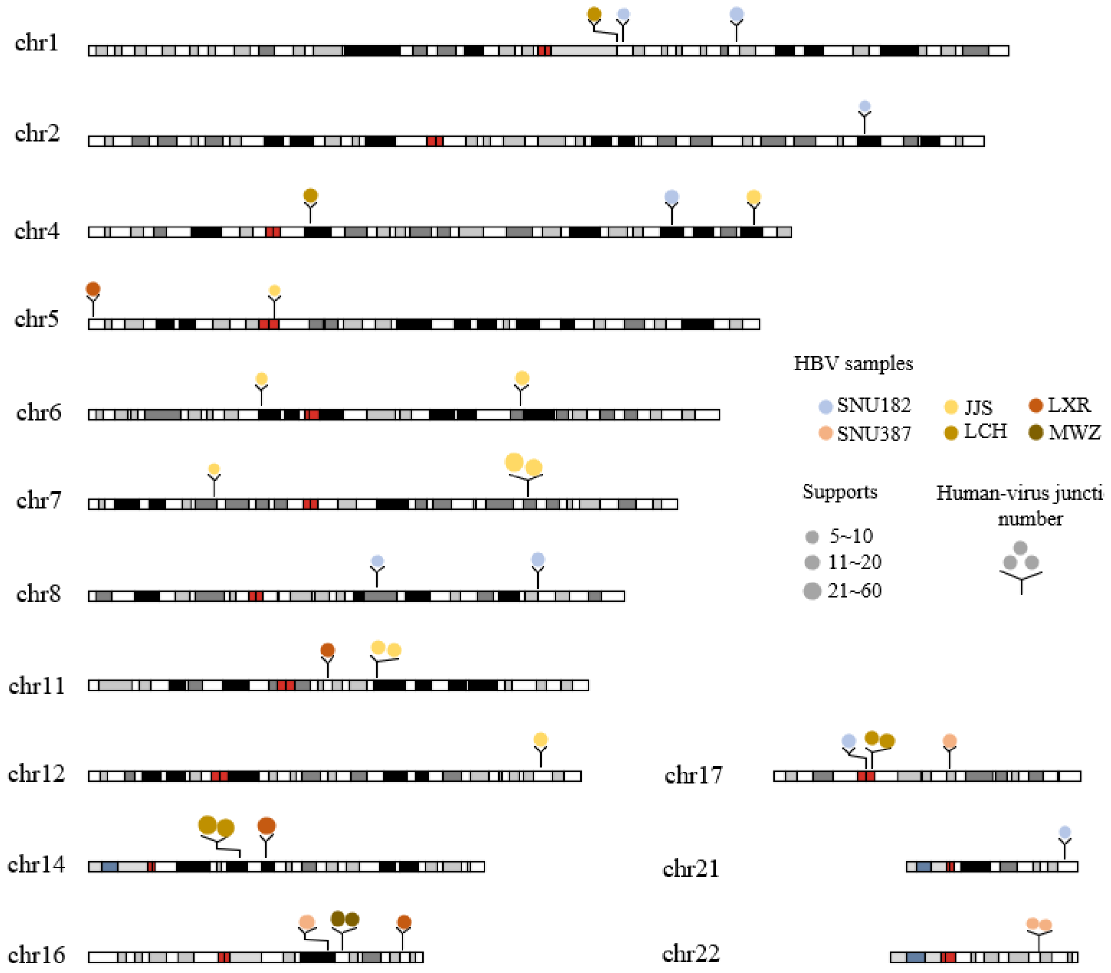

#### Supplementary Figure 2

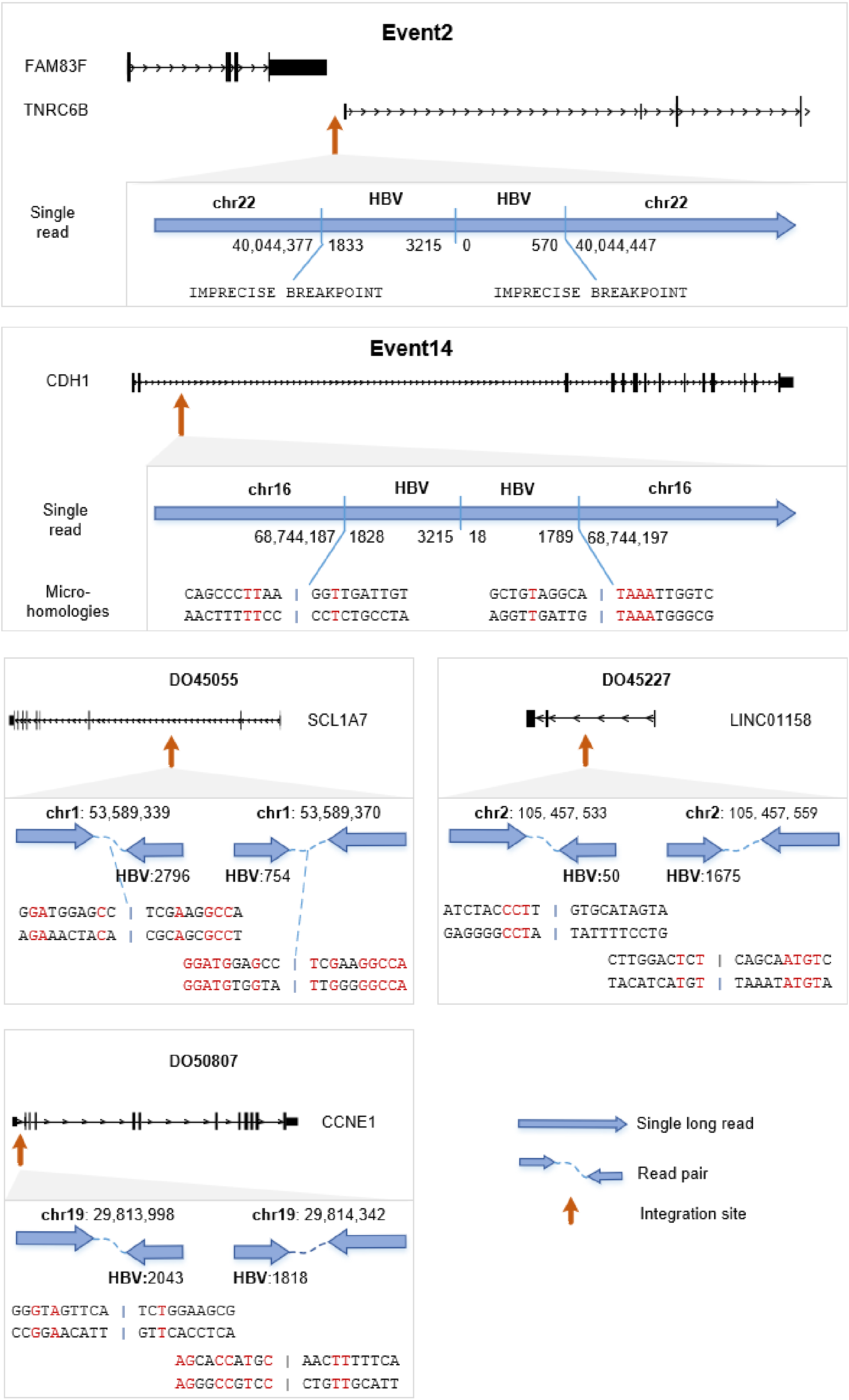

### Supplementary Figure 3

a

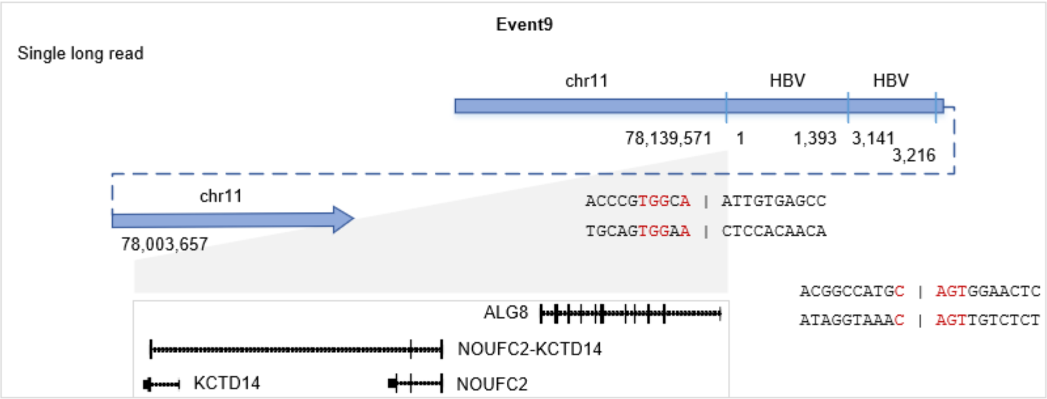


b

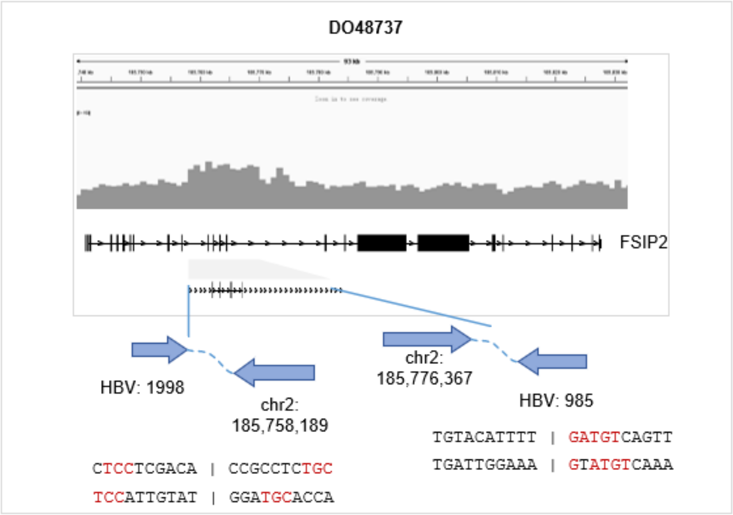

### Supplementary Figure 4

**a**

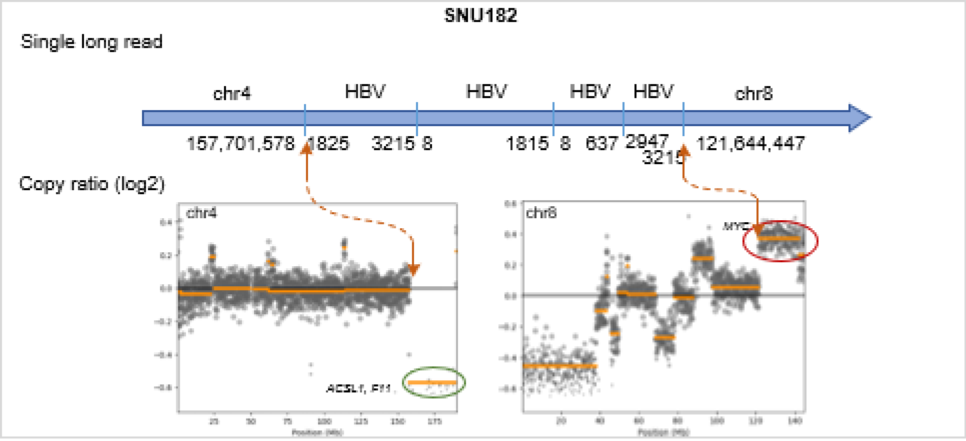


**b**

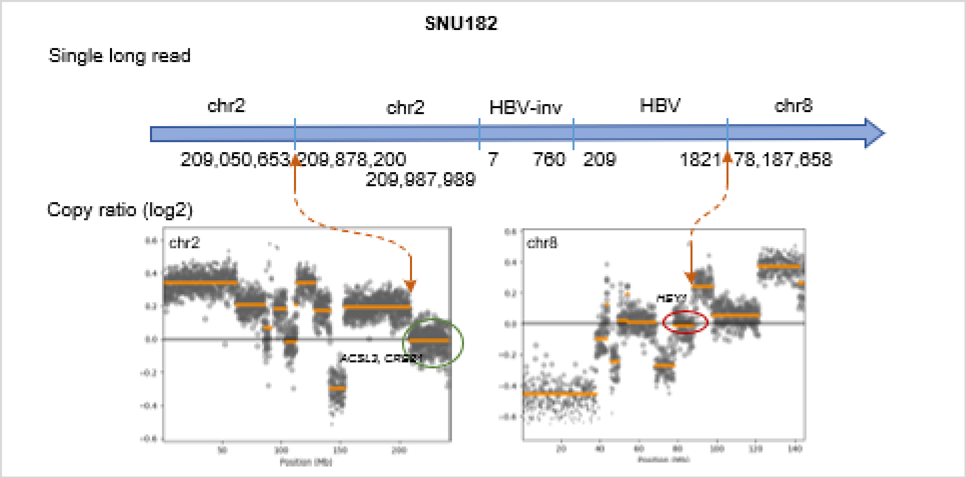

#### Supplementary Figure 5

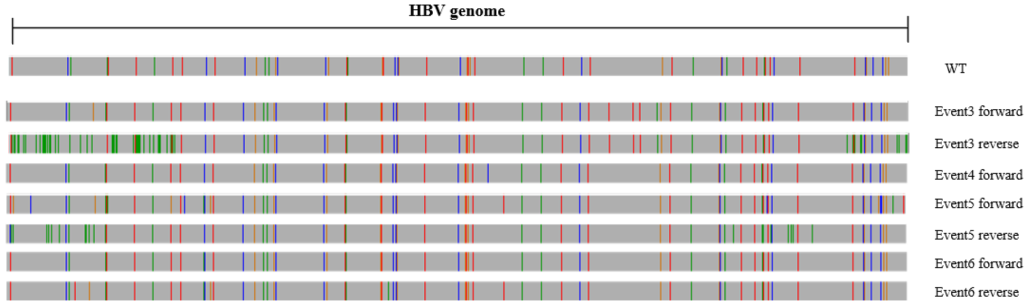


#### Supplementary Figure 6

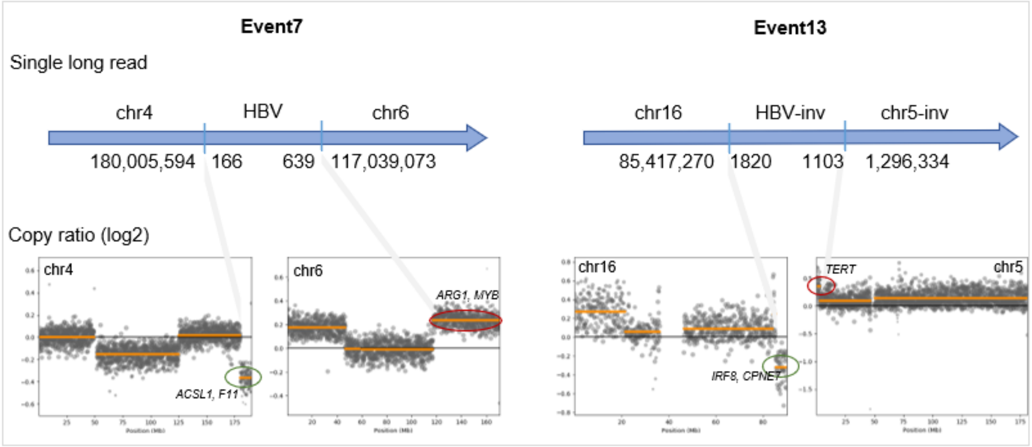


#### Supplementary Figure 7

#### Supplementary Figure 8

**Single long read:** f340e2bc-32da-41ec-b6e4-cd292f2b6d1b, total length: 48,664bp

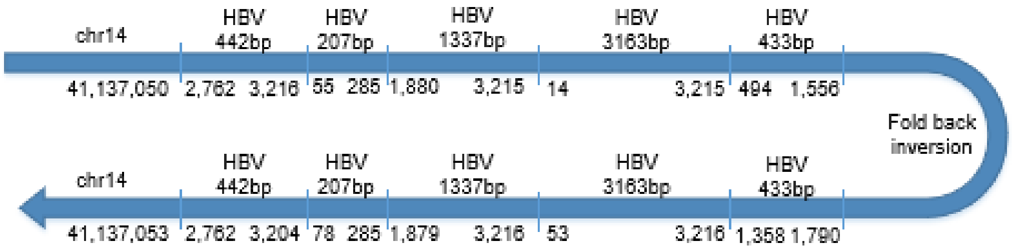


**Dotplot**

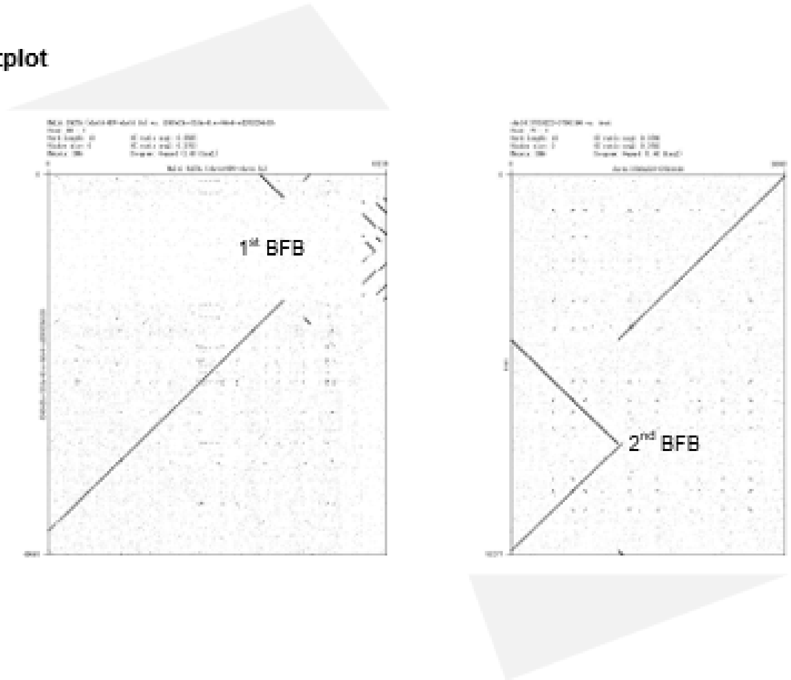


**Single long read:** d0c36459-6427-4b5e-b637-c842f598c984, total length: 26,629bp

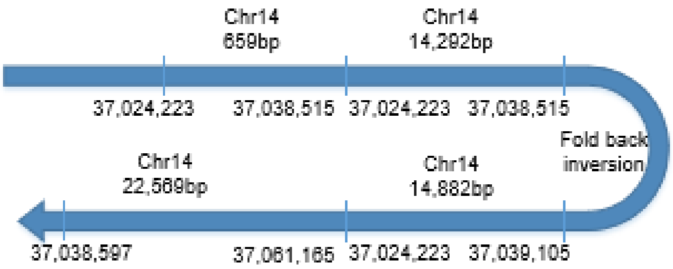

#### Supplementary Figure 9

**Single long read:** f340e2bc-32da-41ec-b6e4-cd292f2b6d1b, total length: 48,664bp

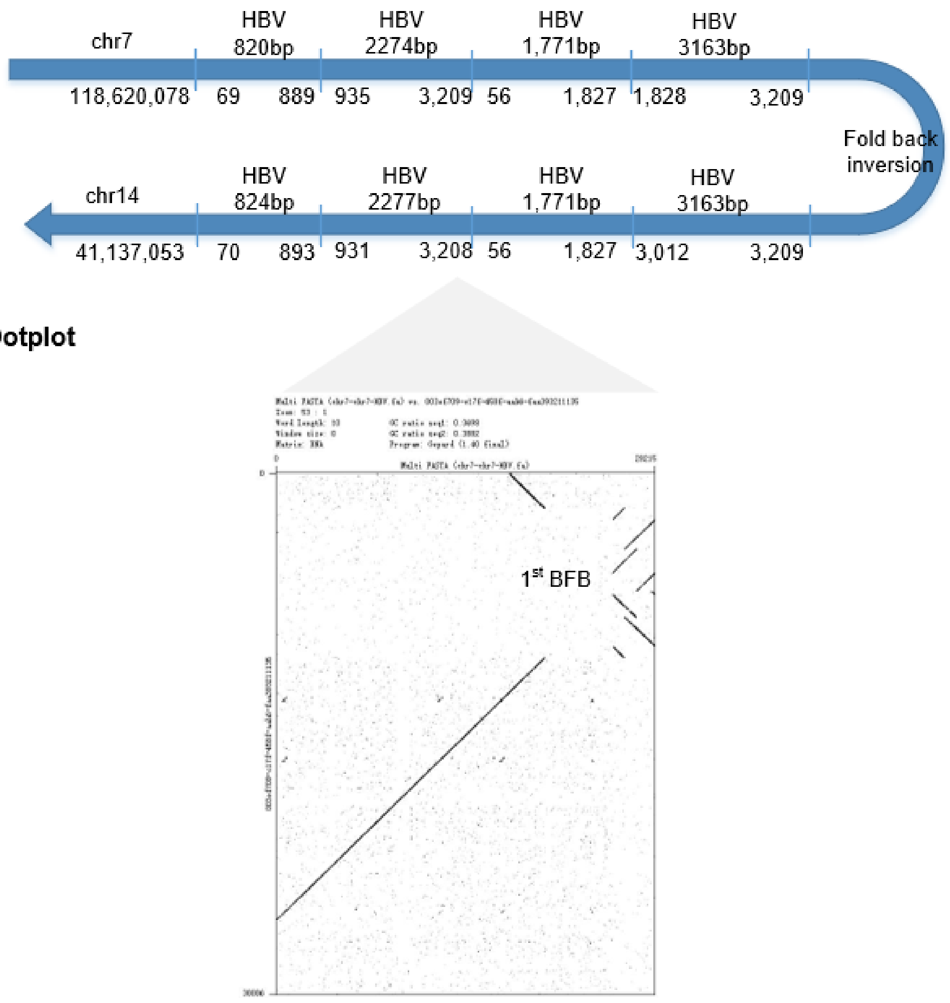

### Supplementary Figure 10

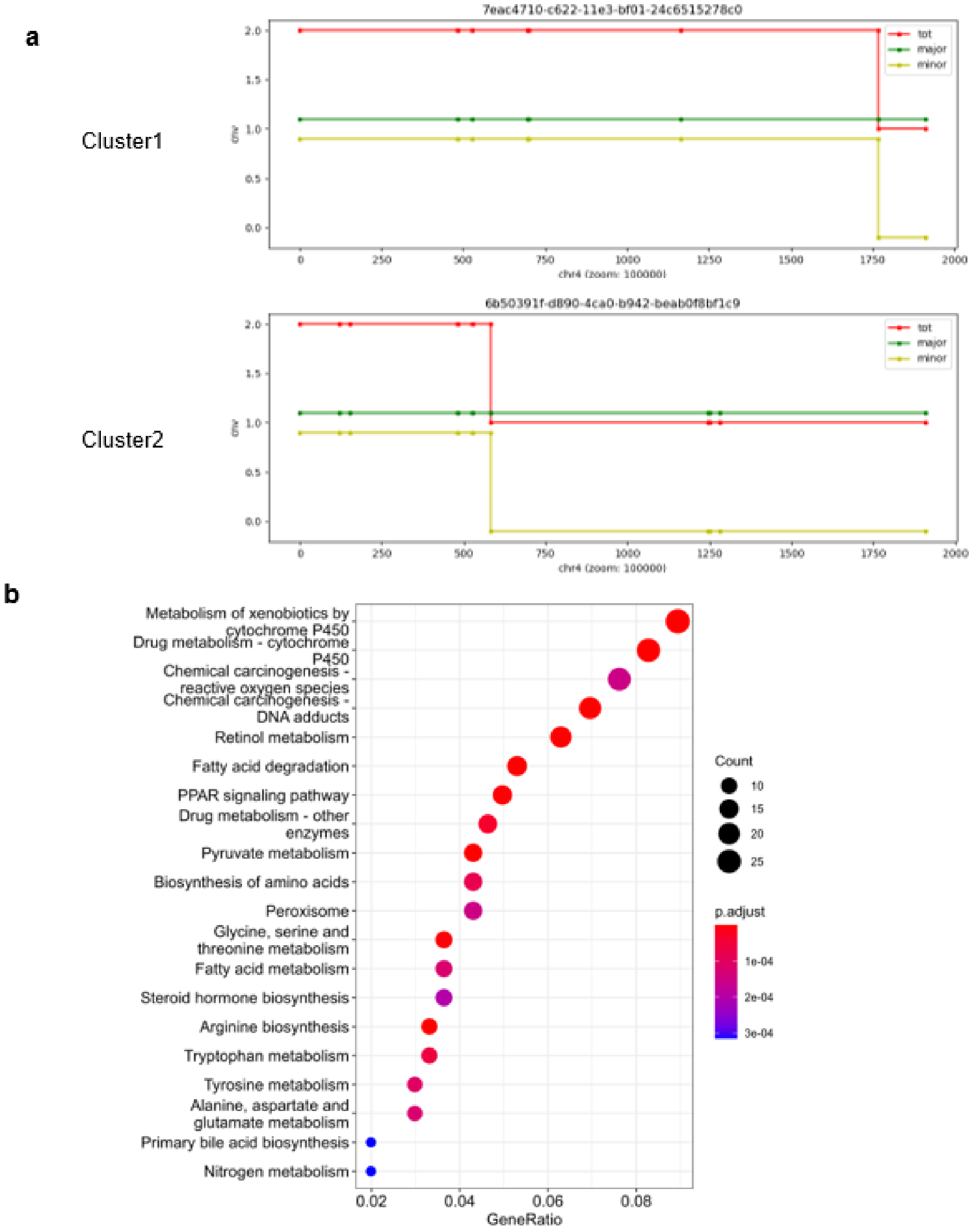
